# HumGut: A comprehensive Human Gut prokaryotic genomes collection filtered by metagenome data

**DOI:** 10.1101/2020.03.25.007666

**Authors:** Pranvera Hiseni, Knut Rudi, Robert C. Wilson, Finn Terje Hegge, Lars Snipen

## Abstract

**Background:** A major bottleneck in the use of metagenome sequencing for human gut microbiome studies has been the lack of a comprehensive genome collection to be used as a reference database. Several recent efforts have been made to re-construct genomes from human gut metagenome data, resulting in a huge increase in the number of relevant genomes. In this work, we aimed to create a collection of the most prevalent healthy human gut prokaryotic genomes, to be used as a reference database, including both MAGs from the human gut and ordinary RefSeq genomes.

**Results:** We screened > 5,700 healthy human gut metagenomes for the containment of > 490,000 publicly available prokaryotic genomes sourced from RefSeq and the recently announced UHGG collection. This resulted in a pool of > 379,000 genomes that were subsequently scored and ranked based on their prevalence in the healthy human metagenomes. The genomes were then clustered at subspecies resolution, and cluster representatives were retained to comprise the HumGut collection. Using the Kraken2 software for classification, we find superior performance in the assignment of metagenomic reads, classifying on average 94.5% of the reads in a metagenome, as opposed to 86% with UHGG and 44% when using standard Kraken2 database. HumGut, half the size of standard Kraken2 database and directly comparable to the UHGG size, outperforms them both.

**Conclusions:** The HumGut collection contains > 30,000 genomes clustered at subspecies resolution and ranked by human gut prevalence. We demonstrate how metagenomes from IBD-patients map equally well to this collection, indicating this reference is relevant also for studies well outside the metagenome reference set used to obtain HumGut. We believe this is a valuable resource in a field in dire need of method standardization. All data and metadata, as well as helpful code, are available at http://arken.nmbu.no/~larssn/humgut/.

## Introduction

Significant efforts have been undertaken to characterize the human gut microbiome, both by microbe isolation and DNA sequencing^1^. A major contribution has also been made by de novo-assembled genomes (Metagenome-Assembled Genomes – MAGs), facilitated by the latest advances in bioinformatics tools^2–6^. As a wrap, a Unified Human Gastrointestinal Genome (UHGG) collection comprised of > 200,000 non-redundant reference genomes was recently announced^7^, marking a major milestone in this field.

These studies have laid a solid foundation, identifying a vast variety of genomes encountered in human guts. However, none of them addresses the global prevalence of genomes within healthy people, i.e. providing information about their frequency of occurrence. This knowledge is essential for setting up a collection of human gut-associated prokaryotic genomes that reflects the worldwide *healthy human gut microbiome*. It is especially important for building custom databases intended to be used for comparative studies in human gastrointestinal microbiome research.

Regionally, studies have shown that the intestinal flora is greatly shaped by the environment^8^ and that its composition can be linked to a range of diseases and disorders^9–12^, thus we are now at a stage where gut microbiota therapeutic interventions are being introduced^13, 14^. However, the lack of a global reference for the intestinal flora in healthy humans represents a bottleneck^15^. This impedes both the understanding of gut microbiota on a worldwide scale and the introduction of large-scale intervention strategies.

The aim of this work was to create a single, comprehensive genome collection of gut microbes associated with healthy humans, called HumGut, as a universal reference for all human gut microbiota studies. We utilized the UHGG collection, mentioned above, along with the NCBI RefSeq genomes. The strategy of building HumGut is outlined in **Figure 1**.

**Figure 1.**
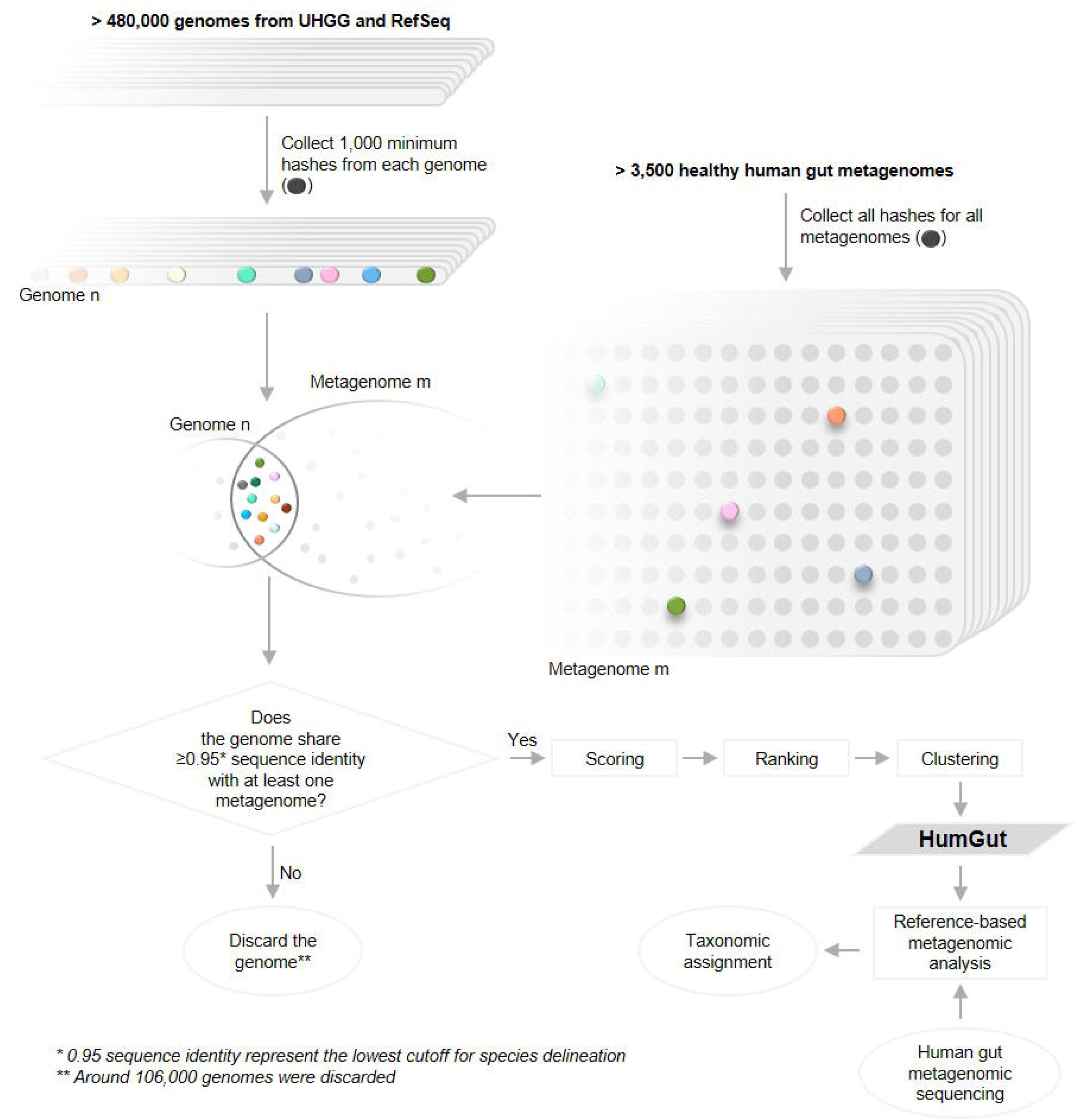
HumGut overview. HumGut represents a collection of genomes and MAGs contained in 3,534 healthy human gut metagenomes. To be considered as contained, a genome shared at least 0.95 sequence identity with at least one of the metagenomes (inferred by the number of shared hashes). The qualified genomes were scored based on the average sequence identity across all the metagenomes. Next, they were ranked based on their scores: the higher the score, the higher the position on the list. Subsequently, the genomes were clustered based on MASH and fastANI distance (D). The top-ranked genome formed a cluster centroid. Around 30,600 clusters were formed applying a D=0.025-threshold. The use of HumGut as a reference set helps the process of taxonomic assignments by drastically reducing the number of unclassified human gut metagenomic reads.

HumGut genomes are ranked by their containment in healthy human gut metagenomes collected worldwide. The most commonly encountered genomes (i.e. top-ranked on the list) were selected as taxa representatives during dereplication, securing thus a list of those most relevant.

While it may seem like a relatively simple concept, this work has only become possible with the recent development of bioinformatics tools that allow the swift screening of publicly available human gut metagenomes for the containment of the ever-growing pool of prokaryotic genomes.

## Results

### Reference metagenomes

More than 5,700 gut metagenome samples collected from healthy people of various ages worldwide were downloaded. These belonged to 72 different BioProjects. To avoid the bias of containing groups of highly similar samples, we computed the MASH distance between metagenomes within each BioProject, then clustered samples with ≥ 95% sequence identity. From each cluster we only kept the medoid sample, resulting thus in a collection of 3,534 healthy human gut metagenomes (**Figure 2a**).

**Figure 2.**
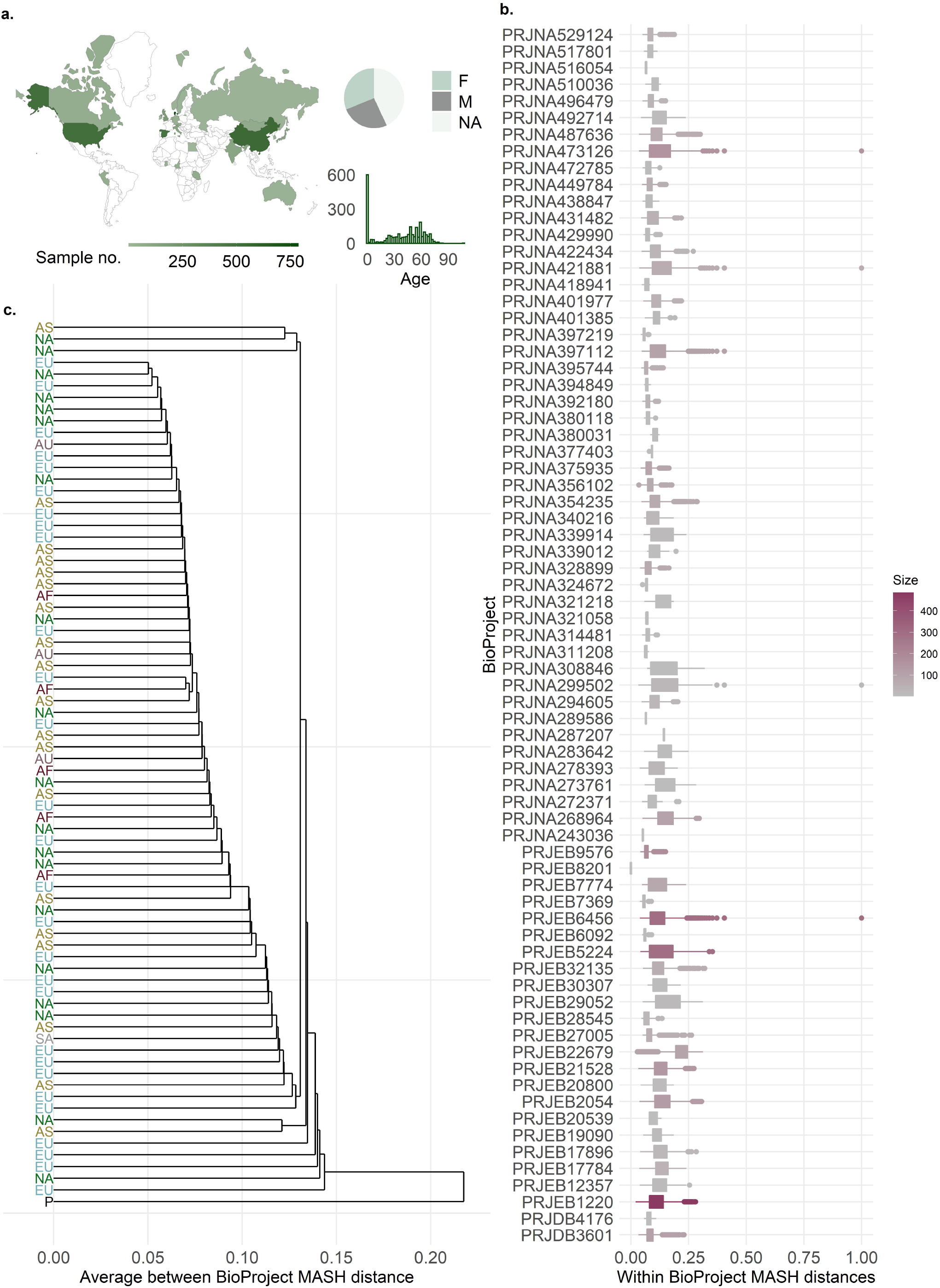
An outline of the metagenomes used in this study. **a**. The geographical, age, and gender distribution of 3,534 metagenomes collected from healthy people. **b**. Boxplots illustrating the distribution of MASH distances between samples within each BioProject. The BioProject accession is used as a label, and the color gradient indicates the size, i.e., the number of samples in each. **c**. Average linkage hierarchical clustering of 72 BioProjects containing healthy samples. BioProjects containing samples from different continents are presented separately. Labels indicate the continent of origin: EU – Europe, AS – Asia, NA – North America, AS – Australia, AF – Africa, SA – South America, and P stands for Primates. Except for the single primate BioProject (BioSample), each BioProject is listed in colored font according to the continent from which it originates. No severe clustering of samples based on origin is detected.

On average, samples within each project shared a 90% sequence identity (*D* = 0.1), indicating a relatively high degree of similarity between one another. There were some outliers, however. Some infant samples (10 belonging to PRJNA473126 project and 1 to PRJEB6456), 10 samples from a project studying the human gut microbiome of vegetarians, vegans, and omnivores (PRJNA421881), and a sample from a study focusing on microbiome diversity among Cheyenne & Arapaho of Oklahoma (PRJNA299502), showed the highest dissimilarity with at least one other sample from the same project (*D* = 1) (**Figure 2b**).

We wanted to see if samples clustered based on their continent of origin (**Figure 2c)**. To do so, we computed the average linkage hierarchical clustering of BioProjects. The distance between two BioProjects is the mean pairwise distance between all their samples. Here, we also included a BioProject containing primate gut metagenome samples (n = 95) as an outgroup against which all human BioProjects were compared. The lowest and highest observed average MASH distances (*D* = 0.05, and *D* = 0.14, respectively) were between two sets of projects stemming from separate continents each, one from Europe and the other from North America. These observations, together with the mixed distribution of BioProjects in the cluster dendrogram, suggested that the clustering of samples did not heavily depend on continent-of-origin. The primate samples were markedly separated from the rest of the tree, showing an average distance of 0.22 from all other BioProjects.

### From genomes to HumGut collection

The majority of genomes stemming from the UHGG collection (99%) and 48% of RefSeq genomes qualified for inclusion in HumGut, resulting thus in a total collection of 381,779 genomes (**Figure 3a**). The qualified genomes were contained within at least one reference metagenome. We inferred the containment by computing sequence identity between genomes and metagenomes using MASH screen, and considered a genome as *contained* when identity was ≥ 0.95.

**Figure 3.**
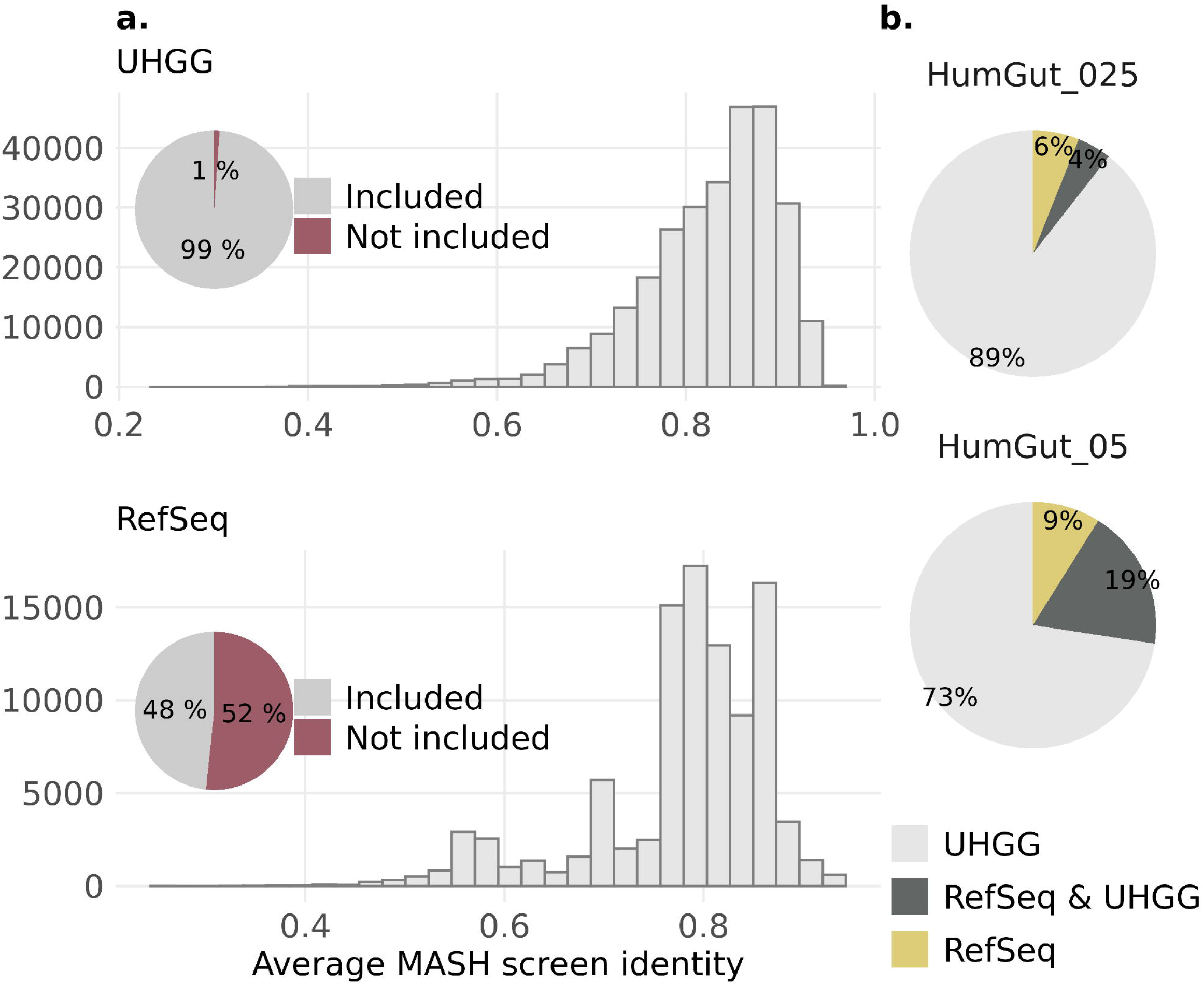
An overview of the genomes used to build HumGut. **a**. The pie charts show the proportion of genomes from each collection (UHGG above, RefSeq below) included in HumGut. To qualify for HumGut inclusion, genomes had to have at least 0.95 MASH screen identity with at least one healthy metagenome, as did most of the UHGG and half of the RefSeq genomes. Histograms show the distribution of the mean identity shared between the qualified genomes and healthy metagenomes. A high average identity means that the qualified genome has been found contained in most of the screened samples **b**. The genome sources for HumGut clusters. The upper pie chart shows data for 30,691 clusters belonging to HumGut_97.5 (genomes grouped based on 97.5% genome sequence identity), the bottom one presents data for 5,170 HumGut_95 clusters (95% sequence identity – species level threshold). The majority of clusters in both HumGut collections are comprised of only UHGG genomes, while 6% and 9% of the clusters consist of only RefSeq genomes (HumGut_97.5, and HumGut_95, respectively).

By applying a rarefaction, we found that the number of new genomes saturated after screening for ca. 1,000 metagenomes, indicating that with > 3,500 metagenomes very few new genomes will be added if screening even more metagenomes from the same population (supplementary material, **Figure S1**).

The most prevalent genomes, i.e., the genomes contained in most metagenomes, belonged to the genus *Bacteroides*, led by *B. vulgatus* (also known as *Phocaeicola vulgatus*), found in more than 70% of samples. It ‘s worth noting that the UHGG collection contained no genome with this species name. The genomes are named as *Bacteroides dorei* instead. We presume that is related to an earlier GTDB database release used for genome taxonomic classifications by Almeida et al (GTDB-Tk v0.3.1; database release 04-RS89)^7^. In the current version of GTDB the species *Phocaeicola vulgatus* is listed.

We performed clustering of genomes based on sequence similarity using the top-ranked genome as a cluster centroid for each iteration. We initially applied an ANI threshold of 97.5% to compile a HumGut collection of highest resolution (HumGut_97.5). This collection resulted in 30,691 genomes with ≥ 50% genome completeness and ≤ 5% contamination. They were all given a GTDB-Tk taxonomic annotation ^16^ as well as an NCBI taxonomy assignment.

These genomes were subsequently clustered further to form a coarser collection at 95% identity, the HumGut_95 with 5,170 genomes. This corresponds roughly to species resolution ^17^.

Looking into genome sources, we found that 9% of HumGut_95 clusters were RefSeq-only genomes (**Figure 3c**). These genomes, 756 in total, clustered into 460 HumGut_95 clusters, belonged to 125 different genera. Most of the genomes (299 in total) belonged to various *Streptococcus* species.

### HumGut genome clusters

Not all species-level clusters were equally diverse, that is not all of them encompassed a similar number of HumGut_97.5 clusters. The majority of HumGut_95 clusters (3,009 out of 5,100) consisted of a single HumGut_97.5 cluster. On the other hand, the most diverse HumGut_95 cluster was one built of 533 different HumGut_97.5 clusters, all named as *Agathobacter rectalis* with GTDB taxonomy (*[Eubacterium] rectale ATCC 33656 with NCBI*). It was followed by a group of 495 subspecies-level clusters consisting of various *Collinsella*-related species names, and a HumGut_95 cluster comprised of 400 different HumGut_97.5 clusters, all GTDB-named as *UBA11524 sp000437595*, and *NCBI*-named as *Faecalibacterium sp. CAG:74*.

Regarding taxonomy, many genomes were not given species names by GTDB, rather they were named after the genus, family, order, or class they belong to. Similarly, the NCBI taxonomic annotations for many genomes resulted in ambiguous names not specific to species, such as for example *uncultured bacterium* or *Firmicutes bacterium*. This contributed greatly in a discrepancy between the total number of species-level clusters (5,170 clusters in HumGut_95) and the total number of distinct cluster names (3,310 GTDB names, 1,716 NCBI names).

There were also many species-level clusters that shared the same species name. This was especially the case with various *Collinsella* clusters, where 81 different GTDB *Collinsella* species gave name to 7 different clusters each, on average. Comparably, 19 NCBI *Collinsella* species were seen in 44 different clusters on average.

### Classifying the metagenome reads

We used the HumGut collection at both resolutions, in addition to the UHGG (species-level collection, containing 4,644 genomes) and the standard Kraken2 database, to classify the metagenomic reads from the 3,534 downloaded samples. On average, there were 56% unclassified reads when using the standard Kraken2 database, while the average dropped substantially when any one of the HumGut or the UHGG collection was utilized (UHGG = 14.1%, Humgut_95 = 11.7%, and HumGut_97.5 = 5.4%, **Figure 4a**).

**Figure 4.**
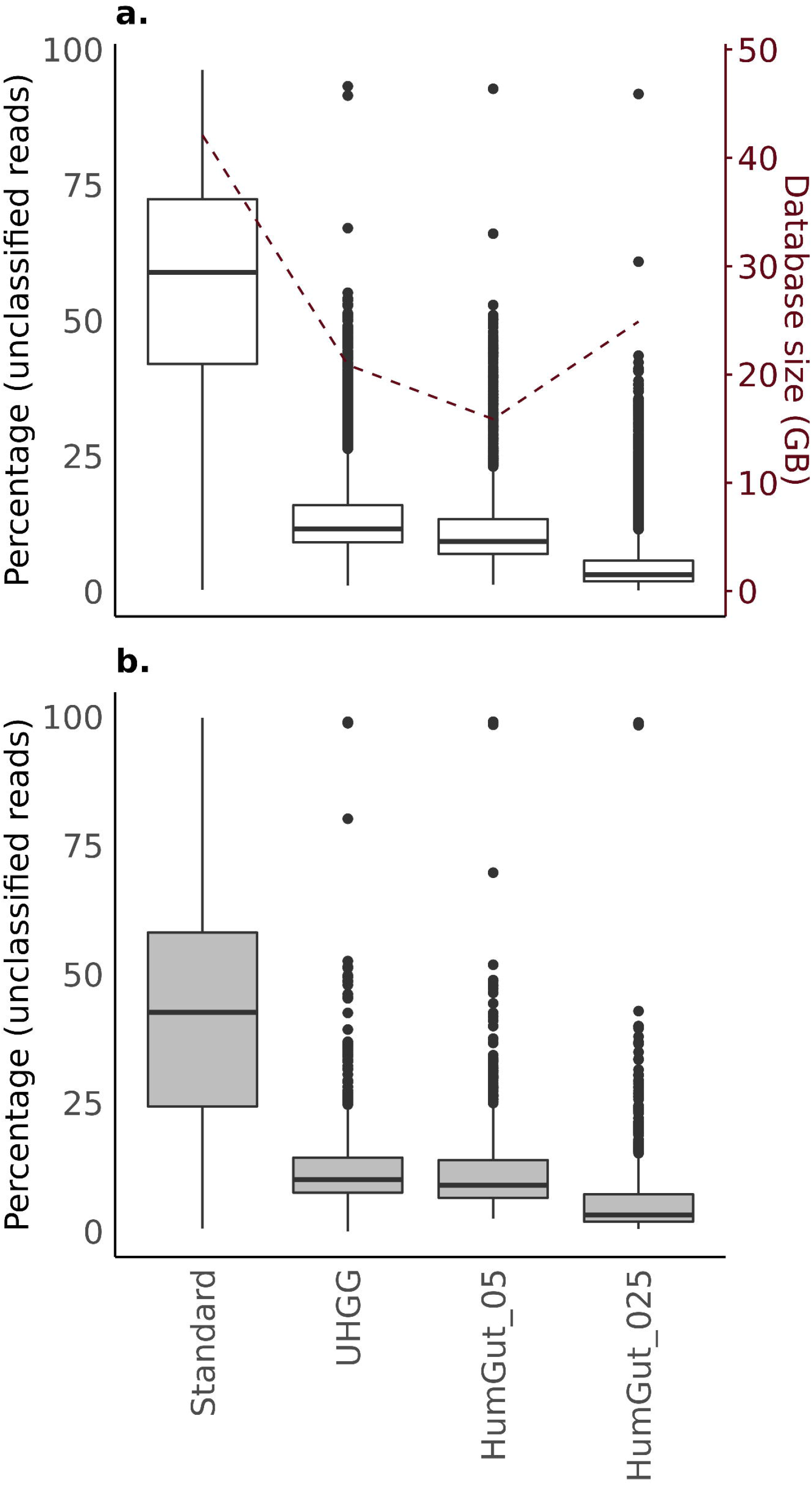
The performance of HumGut versions in comparison to the standard Kraken2 database and UHGG collection. **a**. The boxplot shows the distribution of unclassified reads for the 3,534 analyzed healthy reference metagenome samples. The dashed line represents the k-mer database sizes (right y-axis). Every database version includes standard human genome sequences, in addition to database-specific (sub)sets of bacteria and archaea, and the difference in size is only due to differences in the latter. **b**. The classification of an additional 963 human gut metagenomes, not part of the reference set.

In comparison to the UHGG, both HumGut collections performed better. HumGut_95, a collection of species-level representatives – much like the UHGG collection – classified on average 2.3% more reads than the latter. With HumGut_97.5 as a custom database this increased by 8.7%, marking a significant increase in recognized reads, with an obvious potential for improved classification accuracy.

Both HumGut k-mer databases were smaller than the standard Kraken2 database of k-mers, necessitating reduced computer memory to perform the analyses. The lowest memory was required by the HumGut_95 database (Standard = 42.1 GB, UHGG = 20.9, HumGut_95 = 15.9 GB, HumGut_97.5 = 24.9).

Analysis of an additional 963 gut metagenome samples (collected from people suffering from IBD), not part of the reference set, showed similar results regarding the number of classified reads: 42.3% unclassified reads on average when the Standard database was used, dropping to 6.2% with HumGut_97.5 usage (**Figure 4b**).

### Taxa abundances

We used the KrakenUniq as a means of identifying false positive classifications, and removing them from the Kraken2 reports. We then used the Bracken software on the modified Kraken2 results, to re-estimate species abundance in the classified human gut metagenomes. These tasks were performed using HumGut_97.5 and GTDB taxonomy.

The results showed that, on average, healthy adults contained 202 species, people diagnosed with IBD, 145, and infants, 79 species. The overall species number distribution is presented in **Figure 5a**.

**Figure 5.**
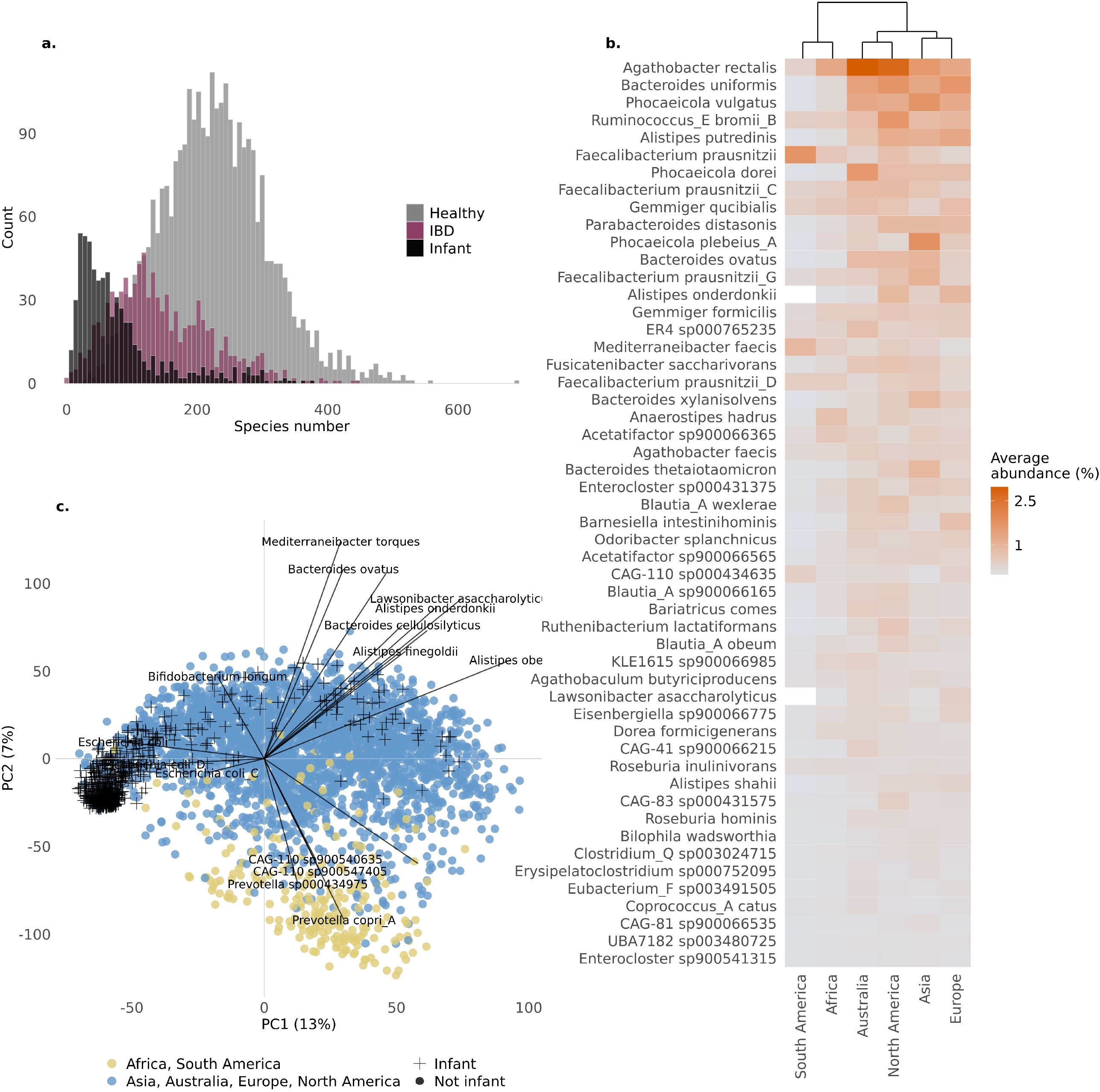
Taxa analysis **a**. Species number distribution between three cohorts: healthy infants, healthy adults, and IBD samples. **b**. Heatplot listing 52 bacterial species found in > 70% healthy adult samples. The tiles are colored based on the average abundance of species in samples from the same continent (Y-axis). On top of the graph, a dendrogram shows the complete linkage between samples from separate continents. **c**. PCA score plot and loading plot. Each dot on the PCA ordination plot represents a metagenome sample. They are colored based on continent of origin. African and South American samples separate more from samples coming from Asia, Australia, Europe, and North America. The superimposed text represents some of the species (loadings) with the strongest effect along the PC2 axis.

In total, 52 species were found present in > 70% of healthy adult samples, led by *Agathobacter rectalis, Blautia_A sp900066165, Bacteroides uniformis, KLE1615 sp900066985, Agathobaculum butyriciproducens* and *Fusicatenibacter saccharivorans*, discovered in > 90% of healthy adult samples, representing a core community of healthy adult human gut microbiota (**Figure 5b**). A complete hierarchical linkage of samples, computed based on the abundance of these top 52 prevalent species, showed that African and South American (*coming exclusively from Peru*) metagenomes were more distant from the rest, while two species were not encountered at all in South American samples (*Alistipes onderdonkii* and *Lawsonibacter assacharolyticus*). In addition, these samples clustered more distantly from the others on a PCA plot (built based on the readcounts from all species), as depicted on **Figure 5c**. The PCA loadings show that *Prevotella* species were more abundant in South American and African samples. In contrast, the *Alistipes* and *Bacteroides* species and lay on the opposite side of the plot, indicating a negative correlation to the former.

Infant samples separated from the adult samples as well. They are represented with crosses instead of dots on the PCA ordination plot, positioned on the leftmost part of the graph along PC1 axis. The loading plot shows that *Escherichia coli* species exercise the largest effect on samples positioned there. The most prevalent bacterium in infants was *Bifidobacterium longum* (68%), followed by *E. coli* (64%).

*Bacteroides vulgatus*, which, after screening the metagenomes using the MASH screen software, was the species of the top scoring genome, was no longer the most prevalent species among healthy human guts when classifying with all HumGut genomes. This was due to a lower diversity among *B. vulgatus* genomes, compared to *Agathobacter rectalis*. The genomes belonging to the former grouped into 2 species-level clusters (*D* = 0.05), while the latter resulted in 16 such groups. It is worth noting that we found the top *B. vulgatus* genome present in 2,536 healthy samples using MASH screen, and we found this species present in 2,537 healthy samples using Kraken2-KrakenUniq-Bracken classification tools. These almost identical results, obtained by two entirely different approaches, increase confidence in the trustworthiness of these findings.

We also investigated the prevalence of species that only had RefSeq as a genome source in our collection. *Streptococcus sanguinis* was found present in 73% of all samples (healthy infants and adults, and IBD), followed by *Flavonifractor sp002161085, Escherichia sp005843885, Streptococcus sp001587175, Pauljensenia sp000466265, Flavonifractor sp002161215, Actinomyces naeslundii, Raoultella terrigena, and Mediterraneibacter sp900120155* (found in 10% - 36% of samples).

## Discussion

The HumGut collection contains 30,691 genomes (HumGut_97.5), with a subset of 5,170 genomes clustered at 95% sequence identity (HumGut_95). The criterion for including a genome in HumGut is its prevalence in healthy human gut metagenomes.

Both HumGut versions showed superior performance in terms of assigned reads compared to the standard Kraken2 database, while demanding far less computational resources, as presented in **Figure 4**. We consider this to be a strong argument in favor of HumGut ‘s comprehensiveness and utility. Classifying a record-high proportion of classified reads per sample, HumGut aids the accuracy of taxonomic classification, which in turn facilitates a next-generation exploration of the human gut microbiome.

The vast majority of UHGG genomes qualified for inclusion in HumGut, as shown in **Figure 3a**. However, in comparison to the UHGG collection, HumGut holds the advantage of containing more relevant human gut prokaryotic genomes in its pool, reflected by the additional RefSeq genomes that showed no sequence similarity with the qualified UHGG genomes, forming separate clusters of 95% sequence identity (**Figure 3b**). An example of its utility is the discovery of *Streptococcus sanguinis* in > 70% of all metagenome samples, which would otherwise be impossible using the UHGG collection as a custom Kraken2 database. Also, the identification of one of the most prevalent species in human guts, *Bacteroides vulgatus*, would have been mistaken for *Bacteroides dorei*.

HumGut sets were built after ranking the genomes based on their prevalence among metagenomes and using the top-ranked ones as cluster representatives. This has ensured that the collections only contain genomes highly relevant to healthy human guts worldwide. Comparing the HumGut_95 collection to the UHGG collection (same resolution) shows that more metagenomic reads are classified for the former. Additionally, its set of unique k-mers is 24% smaller in size than the UHGG. This indicates the UHGG contains a higher genomic diversity, requiring memory which is not really needed for successful read classification. These are rare genomes found in the occasional human gut metagenome, but with low prevalence.

HumGut can serve as a global reference for bacteria inhabiting the gut of healthy humans, highlighting its importance for future gut microbiome studies and is available for download (http://arken.nmbu.no/~larssn/humgut/).

Our analysis showed that the diversity of gut samples across the world is not profoundly affected by geography (**Figure 2**); therefore, having a global genome collection like HumGut is reasonable. We found 50 bacterial species present in more than 70% of the samples, regardless of the country of origin. This group of species, led by *Agathobacter rectalis*, represents what we think is the core human gut bacterial community (**Figure 5b**). We must, however, cautiously refer to A. *rectalis* as the most prevalent/abundant species found in human gut samples. That because we found this species to be highly diverse in sequence identity. In our collection, we have 16 different species-level clusters, and more than 530 subspecies-level clusters with this name.

We discovered that, on average, healthy adults contain around 60 bacterial species more than IBD subjects, and around 120 species more than healthy infants (**Figure 5a**). A low microbiome complexity among the latter two groups is well documented in literature^18–22^.

Although we found a great homogeneity of top prevalent species among healthy adults, our analysis showed that samples originating from Africa and South America were rich in *Prevotella* and poor in *Bacteroides*, which made them cluster in our principal component analysis, as depicted in **Figure 5c**. A *Prevotella* – *Bacteroides* antagonism and their correlation to lifestyle and diet have long been described in literature^23, 24^. Our results are, therefore, consistent with these findings.

We have demonstrated that HumGut is useful in research that goes beyond studying healthy subjects, exemplified by the equally high number of classified metagenomic reads collected from IBD subjects.

A challenge that remains is the nomenclature of species in our genome collection. There is a profound inconsistency between the total number of species-level clusters and the total number of names annotating them (a ratio of 1.5:1 with GDTB-based annotation, and 3:1 with NCBI names). We believe that as long as not all names reflect species individuality, it will be difficult to truly explore the composition differences between various cohorts, in addition to posing a challenge in studies linking functions to species. On our website, we have prepared files for building a custom Kraken2 database where all HumGut clusters also have been given artificial ‘taxonomy IDs, ‘ making it possible to classify to clusters instead of taxa. We note that the decision regarding which version the HumGut collection to employ depends on users ‘ computational resources as well as the level of taxonomic resolution required.

As future work, we will also extend our approach to more disease-associated genomes and metagenomes, in addition to screening for gut genomes that will expectedly be published in the future.

## Conclusion

We believe that by using HumGut as a reference, the scientific community will be one step closer to method standardization sorely needed in the field of human gut microbiome analysis, and that the discovery of potential microbiome markers will be facilitated with higher certainty in less time and computational resources.

## Methods

### Human gut reference metagenomes

A set of publicly available human gut metagenome samples were collected and used for ranking all genomes in the search for human gut relevant ones. A text search for all human gut microbiome samples at the Sequence Read Archive (NCBI/SRA, https://www.ncbi.nlm.nih.gov/sra) was performed. The list of hits was manually curated, keeping only samples with > 1,000,000 reads and annotated as healthy individuals. NCBI/BioProject accessions of these projects were used to locate the same data in the European Nucleotide Archive (EMBL-EBI/ENA, https://www.ebi.ac.uk/ena)” www.ebi.ac.uk/ena), from which all samples were downloaded as compressed fastq-files, using the Aspera download system (https://www.ibm.com/products/aspera). This resulted in 5,737 healthy metagenomes (samples) covering 74 BioProjects. For many BioProjects, some samples tended to be very similar to each other, presumably due to samples collected from individuals sharing the same lifestyle, geographical sub-population, genetics, or other factors that may affect the human gut microbiome. To avoid too much bias in the direction of such heavily sampled sub-populations, samples from the same BioProject were clustered. From each metagenome sample, a MinHash sketch of 10,000 k-mers was computed using the MASH software^25^, discarding singleton k-mers (21-mers). Based on these sketches the MASH distances between all pairs of samples were calculated. A MASH distance close to zero means two samples are very similar, sharing most of their k-mers. Next, hierarchical clustering with complete linkage was computed, and samples were partitioned at a 0.05 distance threshold, resulting in clusters with ‘diameters ‘ no larger than this chosen threshold. The medoid sample from each cluster, i.e., the one with the minimum sum of distances to all members of the cluster, was retained as the reference sample representing its cluster. This resulted in the set of 3,534 healthy metagenome samples. Below we refer to this collection as *MetHealthy*.

The same procedure was utilized to collect another set of metagenomes from patients diagnosed with Irritable Bowel Disease (Ulcerative Colitis, or Crohn ‘s Disease). From initially 2,064 metagenomes the clustering resulted in a collection of 963 metagenomes covering 14 BioProjects. This is the *MetIBD* collection. Finally, a set of 95 samples containing gut metagenome data from primates was collected and used as an outgroup in a comparison of the human gut metagenomes. The metagenomes ‘ metadata is included in the **supplementary Table 1**.

### Genome collections

The main source was the recently published Unified Human Gut Genomes (UHGG) collection, containing 286,997 genomes exclusively related to human guts:http://ftp.ebi.ac.uk/pub/databases/metagenomics/mgnify_genomes/human-gut/v1.0/all_genomes/. The other source was NCBI/Genome, the RefSeq repository at ftp://ftp.ncbi.nlm.nih.gov/genomes/refseq/bacteria/ and ftp://ftp.ncbi.nlm.nih.gov/genomes/refseq/archaea/. At the time of writing, ∼204,000 genomes were downloaded from this site.

Metadata about the genomes considered and qualified for HumGut are presented on supplementary **Table 2**.

### Genome ranking

Only metagenomes collected from healthy individuals, *MetHealthy*, were used in this step. For all genomes, the MASH software was again used to compute sketches of 1,000 k-mers, including singletons^26^. The MASH screen compares the sketched genome hashes to all hashes of a metagenome, and, based on the shared number of them, estimates the genome sequence identity *I* to the metagenome. Given that *I = 0*.*95* (95% identity) is regarded as a species delineation for whole-genome comparisons^17^, it was used as a soft threshold to determine if a genome was contained in a metagenome. Genomes meeting this threshold for at least one of the *MetHealthy* metagenomes were qualified for further processing. Then the average *I*-value across all *MetHealthy* metagenomes was computed for each genome, and this prevalence-score was used to rank them. The genome with the highest prevalence-score was considered the most prevalent among the *MetHealthy* samples, and thereby the best candidate to be found in any healthy human gut. This resulted in a list of genomes ranked by their prevalence in healthy human guts.

### Genome clustering

Many ranked genomes were very similar, some even identical. Due to errors introduced in sequencing and genome assembly, it made sense to group genomes and use one member from each group as a representative genome. Even without any technical errors, a lower meaningful resolution in terms of whole genome differences was expected, i.e., genomes differing in only a small fraction of their bases should be considered identical.

The clustering of the genomes was performed in two steps, like the procedure used in the dRep software^27^, but in a greedy way based on the ranking of the genomes. The huge number of genomes (hundreds of thousands) made it extremely computationally expensive to compute all-versus-all distances. The greedy algorithm starts by using the top ranked genome as a cluster centroid, and then assigns all other genomes to the same cluster if they are within a chosen distance *D* from this centroid. Next, these clustered genomes are removed from the list, and the procedure is repeated, always using the top ranked genome as centroid.

The whole-genome distance between the centroid and all other genomes was computed by the fastANI software^17^. However, despite its name, these computations are slow in comparison to the ones obtained by the MASH software. The latter is, however, less accurate, especially for fragmented genomes. Thus, we used MASH-distances to make a first filtering of genomes for each centroid, only computing fastANI distances for those who were close enough to have a reasonable chance of belonging to the same cluster. For a given fastANI distance threshold *D* we first used a MASH distance threshold *D*_mash_ *>> D* to reduce the search space. In supplementary material, **Figure S2**, we show some results guiding the choice of *D*_*mash*_ for a given *D*.

A distance threshold of D = 0.05 is regarded as a rough estimate of a species, i.e. all genomes within a species are within this fastANI distance from each other^16, 17^. This threshold was also used to arrive at the 4,644 genomes extracted from the UHGG collection and presented at the MGnify website. However, given shotgun data, a larger resolution should be possible, at least for some taxa. For this reason, we started out with a threshold D = 0.025, i.e. half the “species radius.” An even higher resolution was tested (*D = 0*.*01*), but the computational burden increases vastly as we approach 100% identity between genomes. It is also our experience that genomes more than ∼98% identical are very difficult to separate, given today ‘s sequencing technologies^28^. However, the genomes found at D = 0.025 (HumGut_97.5) were also again clustered at D = 0.05 (HumGut_95) giving two resolutions of the genome collection.

The taxonomic annotation of the genomes was performed with GTDB toolkit (GTDB-Tk, release 05-RS95, https://github.com/Ecogenomics/GTDBTk)^16^, but in the genome metadata tables we provide on our website, we made efforts to list the corresponding NCBI Taxonomy names and ID ‘s for all genomes.

All UHGG genomes were already checked for completeness and contamination^7^. The completeness and contamination of RefSeq genomes was performed using CheckM (https://ecogenomics.github.io/CheckM/)^29^. The handful genomes not having > 50% completeness and < 5% contamination were discarded.

### Metagenome classifications

The Kraken2 software was used for classifying reads from the metagenome samples^30^. To see the effects of using a different database, the standard Kraken2-database was compared by custom databases built from the 4,644 UHGG genomes at the MGnify website as well as our HumGut collections. In all custom databases, the standard Kraken2 library for the human genome was also included, since host contamination is quite normal in metagenome data. All classifications were performed using default settings in Kraken2.

Since Kraken2, like most other software for taxonomic classification, uses the Lowest Common Ancestor (LCA) approach, many reads are assigned to ranks high up in the taxonomy. The Bracken software^31^ has been designed to re-estimate the abundances at some fixed rank, by distributing reads from higher ranks into the lower rank, based on conditional probabilities estimated from the database content. A Bracken database (100-mers) was created for HumGut_97.5 database and used to re-estimate all abundances at the species rank.

If counting all listed taxa, regardless of low readcounts, the Kraken2 is known to produce many false positives^32^, i.e. list taxa as present when they are in fact not. The KrakenUniq software has been developed to handle this problem^32^. We ran it to classify the metagenome reads for both healthy and IBD metagenomes. The overall results from both Kraken2 and KrakenUniq tools were similar, but KrakenUniq also reports the number of unique k-mers in each genome covered by the reads. On the other hand, only Kraken2 reports are compatible for running Bracken software. Since we were interested in both - that is finding the true positive identifications, and their estimated abundances - we combined the two approaches. For each sample, we found from the KrakenUniq report a k-mer count threshold, following the authors recommendations (2,000 unique k-mers per 1,000,000 sequencing reads depth)^32^. Taxa falling below this threshold were given zero read counts in the corresponding modified Kraken2 reports. We then ran Bracken on these modified Kraken2 reports.

A Principal Component Analysis was conducted on the matrix of species readcounts for all metagenome samples, after the following transformation: a pseudo-count of 10 was added to all species before using Aitchison ‘s centered log-ratio transform^33, 34^ to remove the unit-sum constraint otherwise affecting a PCA of such data.

## Declarations

### Ethics approval and consent to participate

Not applicable

### Consent for publication

Not applicable

### Availability of data and material

The HumGut genome collection and all associated metadata can be found at http://arken.nmbu.no/~larssn/humgut/. This also includes files and recipes for building Kraken2 databases using these genomes.

## Acknowledgements

Not applicable

## Author contributions

LS conceived the study. LS and PH worked out the technical aspects of the paper. All authors discussed and interpreted the results. PH wrote the article with equal input from all authors.

## Figure legend

**Figure S1.**
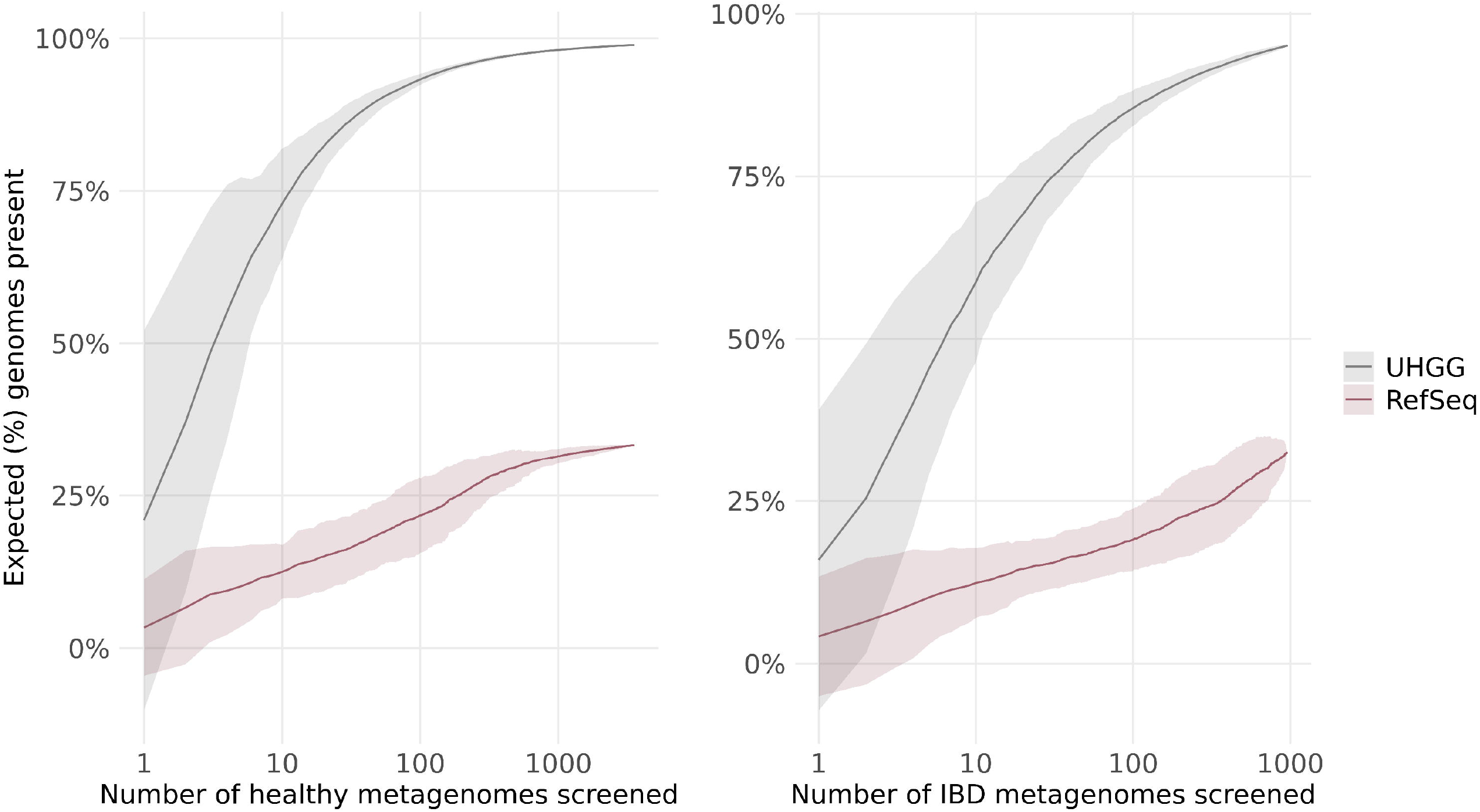
Rarefaction curves for healthy (left panel) and IBD metagenomes (right panel), showing that the number of new expected genomes flattens after screening ca. 1,000 metagenomes.

**Figure S2.**
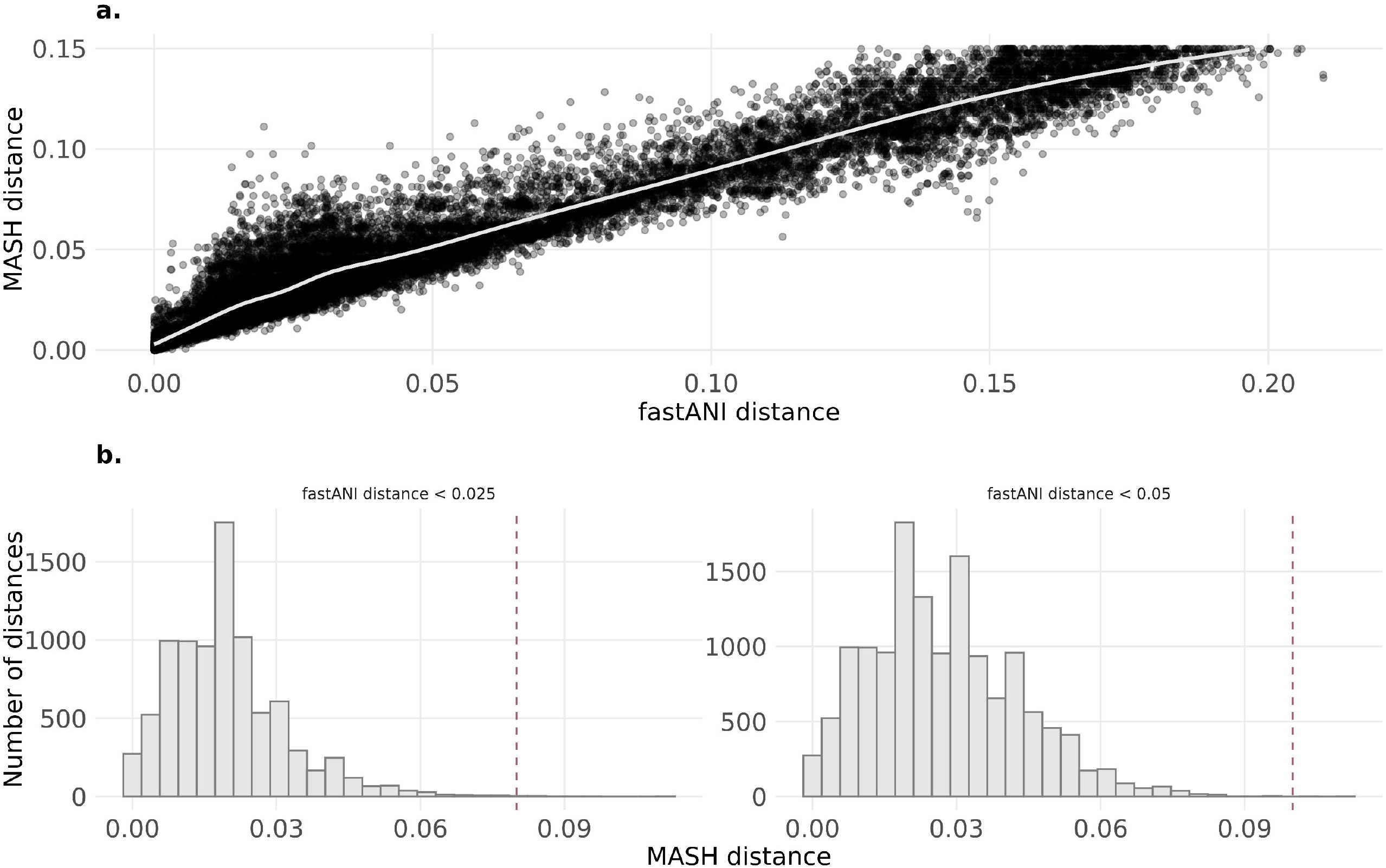
MASH and fastANI distances. **a**. A plot of ca. 20,000 genome distances computed with both fastANI (x-axis) and MASH (y-axis). fastANI distances tend to be a little smaller than MASH distances, they however have a substantial variance. **b**. The rationale behind using 0.08, and 0.1 MASH distance thresholds (vertical dashed lines) for HumGut clustering algorithm. The vast majority of fastANI distances < 0.025 have a MASH distance < 0.08 and genomes with fastANI < 0.05 have a MASH distance < 0.1. When clustering, the distance between all genomes was first computed using MASH, then only genomes with distances below the abovementioned thresholds were included to speed up fastANI computations.

## Literature

1. Methé, B.A., et al., A framework for human microbiome research. Nature, 2012. 486(7402): p. 215–221.

2. Forster, S.C., et al., A human gut bacterial genome and culture collection for improved metagenomic analyses. Nature Biotechnology, 2019. 37(2): p. 186–192.

3. Almeida, A., et al., A new genomic blueprint of the human gut microbiota. Nature, 2019. 568(7753): p. 499–504.

4. Zou, Y., et al., 1,520 reference genomes from cultivated human gut bacteria enable functional microbiome analyses. Nature Biotechnology, 2019. 37(2): p. 179–185.

5. Nayfach, S., et al., New insights from uncultivated genomes of the global human gut microbiome. Nature, 2019. 568(7753): p. 505–510.

6. Pasolli, E., et al., Extensive Unexplored Human Microbiome Diversity Revealed by Over 150,000 Genomes from Metagenomes Spanning Age, Geography, and Lifestyle. Cell, 2019. 176(3): p. 649-662.e20.

7. Almeida, A., et al., A unified catalog of 204,938 reference genomes from the human gut microbiome. Nature Biotechnology, 2020.

8. Rothschild, D., et al., Environment dominates over host genetics in shaping human gut microbiota. Nature, 2018. 555(7695): p. 210–215.

9. Lozupone, C.A., et al., Diversity, stability and resilience of the human gut microbiota. Nature, 2012. 489(7415): p. 220–230.

10. Shin, N.-R., T.W. Whon, and J.-W. Bae, Proteobacteria: microbial signature of dysbiosis in gut microbiota. Trends in Biotechnology, 2015. 33(9): p. 496–503.

11. Halfvarson, J., et al., Dynamics of the human gut microbiome in inflammatory bowel disease. Nature Microbiology, 2017. 2(5): p. 17004.

12. Rajilić–Stojanović, M., et al., Global and Deep Molecular Analysis of Microbiota Signatures in Fecal Samples From Patients With Irritable Bowel Syndrome. Gastroenterology, 2011. 141(5): p. 1792–1801.

13. Cotillard, A., et al., Dietary intervention impact on gut microbial gene richness. Nature, 2013. 500(7464): p. 585–588.

14. Wallace, T.C., et al., Human gut microbiota and its relationship to health and disease. Nutrition Reviews, 2011. 69(7): p. 392–403.

15. McBurney, M.I., et al., Establishing What Constitutes a Healthy Human Gut Microbiome: State of the Science, Regulatory Considerations, and Future Directions. The Journal of nutrition, 2019. 149(11): p. 1882–1895.

16. Chaumeil, P.-A., et al., GTDB-Tk: a toolkit to classify genomes with the Genome Taxonomy Database. Bioinformatics, 2019. 36(6): p. 1925–1927.

17. Jain, C., et al., High throughput ANI analysis of 90K prokaryotic genomes reveals clear species boundaries. Nature communications, 2018. 9(1): p. 5114.

18. Manichanh, C., et al., Reduced diversity of faecal microbiota in Crohn ‘s disease revealed by a metagenomic approach. 2006. 55(2): p. 205–211.

19. Manichanh, C., et al., The gut microbiota in IBD. Nature Reviews Gastroenterology & Hepatology, 2012. 9(10): p. 599–608.

20. Matsuoka, K. and T. Kanai, The gut microbiota and inflammatory bowel disease. Seminars in Immunopathology, 2015. 37(1): p. 47–55.

21. RodrÍguez, J.M., et al., The composition of the gut microbiota throughout life, with an emphasis on early life. Microbial Ecology in Health and Disease, 2015. 26(1): p. 26050.

22. Moore, R.E. and S.D. Townsend, Temporal development of the infant gut microbiome. 2019. 9(9): p. 190128.

23. Gorvitovskaia, A., S.P. Holmes, and S.M. Huse, Interpreting Prevotella and Bacteroides as biomarkers of diet and lifestyle. Microbiome, 2016. 4(1): p. 15.

24. Hjorth, M.F., et al., Prevotella-to-Bacteroides ratio predicts body weight and fat loss success on 24-week diets varying in macronutrient composition and dietary fiber: results from a posthoc analysis. International Journal of Obesity, 2019. 43(1): p. 149–157.

25. Ondov, B.D., et al., Mash: fast genome and metagenome distance estimation using MinHash. Genome biology, 2016. 17(1): p. 132.

26. Ondov, B.D., et al., Mash Screen: High-throughput sequence containment estimation for genome discovery. BioRxiv, 2019: p. 557314.

27. Olm, M.R., et al., dRep: a tool for fast and accurate genomic comparisons that enables improved genome recovery from metagenomes through de-replication. The ISME Journal, 2017. 11(12): p. 2864–2868.

28. Snipen, L., et al., Reduced Metagenome Sequencing for strain-resolution taxonomic profiles. Microbiome, 2021.

29. Parks, D.H., et al., CheckM: assessing the quality of microbial genomes recovered from isolates, single cells, and metagenomes. Genome Res, 2015. 25(7): p. 1043–55.

30. Wood, D.E., J. Lu, and B. Langmead, Improved metagenomic analysis with Kraken 2. Genome biology, 2019. 20(1): p. 257.

31. Lu, J., et al., Bracken: estimating species abundance in metagenomics data. PeerJ Computer Science, 2017. 3: p. e104.

32. Breitwieser, F.P., D.N. Baker, and S.L. Salzberg, KrakenUniq: confident and fast metagenomics classification using unique k-mer counts. Genome Biology, 2018. 19(1): p. 198.

33. Weiss, S., et al., Normalization and microbial differential abundance strategies depend upon data characteristics. Microbiome, 2017. 5(1): p. 27.

34. Aitchison, J., The statistical analysis of compositional data. Journal of the Royal Statistical Society: Series B (Methodological), 1982. 44(2): p. 139–160.

